# Effects of acclimation time and epigenetic mechanisms on growth of *Neurospora* in fluctuating environments

**DOI:** 10.1101/234971

**Authors:** Ilkka Kronholm, Tarmo Ketola

## Abstract

Reaction norms or tolerance curves have often been used to predict how organisms deal with fluctuating environments. A potential drawback is that reaction norms measured in different constant environments may not capture all aspects of organismal responses to fluctuating environments. We examined growth of the filamentous fungus *Neurospora crassa* in fluctuating temperatures and tested if growth in fluctuating temperatures can be explained simply by growth in different constant temperatures or if more complex models are needed. In addition, as previous studies on fluctuating environments have revealed that past temperatures that organisms have experienced can affect their response to current temperature, we tested the roles of different epigenetic mechanisms in response to fluctuating environments using different mutants. We found that growth of *Neurospora* can be predicted in fluctuating temperatures to some extent if acclimation times are taken into account in the model. Interestingly, while fluctuating environments have been linked with epigenetic responses we found only some evidence of involvement of epigenetic mechanisms on tolerating fluctuating temperatures. Mutants which lacked H3K4 or H3K36 methylation had slightly impaired response to temperature fluctuations, in addition the H3K4 methylation mutant and a mutant in the RNA interference pathway had altered acclimation times.

## Introduction

Phenotypic plasticity is the ability of organisms to change or develop different phenotypes in response to environmental changes (DeWitt and Scheiner, 2004). We can describe phenotypic plasticity using reaction norms, where the phenotypic value of a particular trait is a function of an environmental variable. For example, the growth of a single genotype in different temperatures. Natural selection can act on plasticity (Scheiner, 2002; Nussey *et al*., 2005), and plasticity is important in life-history theory (Day and Rowe, 2002), and sexual selection theory (Ingleby *et al*., 2010). Plasticity can also influence evolutionary dynamics, when a population adapts to a new environment (Lande, 2009; Chevin *et al*., 2010). In terms of practical applications, reaction norms for temperature, also called thermal performance curves, have been used in predicting the effects of climate change on extinction risk in ectotherm populations (Deutsch *et al*., 2008; Kingsolver *et al*., 2013; Vasseur *et al*., 2014; Levy *et al*., 2015).

Reaction norms are a useful tool, but there exists a significant complication to their practical application. Reaction norms are often measured in the laboratory in constant environmental conditions. In the wild most environments rarely stay constant for long: day and night cycles, seasons, and daily changes in weather all cause the environment to change constantly. This may be a significant drawback for the usefulness of reaction norms, as recently there have been reports that reaction norms measured in constant environments do not predict the performance of organisms in fluctuating environments (Kingsolver *et al*., 2009; Niehaus *et al*., 2012; Rezende *et al*., 2014; Ketola *et al*., 2014; Ketola and Saarinen, 2015; Kingsolver *et al*., 2015; Saarinen *et al*., 2018). Ketola and Kristensen (2017) and Sinclair *et al*. (2016) list the key complications of why reaction norms measured in the lab may not predict performance in natural environments: performance depends on the current temperature but also on temperatures experienced in the past (Schulte *et al*., 2011; Rezende *et al*., 2014); the time an organism has experienced the current temperature may influence acclimation, as acclimation to a given temperature may not be instantaneous (Johnston and Dunn, 1987). The duration of exposure or the frequency of environmental fluctuations may also be important (Ketola and Kristensen, 2017). For example, the benefits of inducible plasticity could be diminished if response time for phenotypic adjustment is slower than a new alteration in the environment (Padilla and Adolph, 1996).

Mechanisms of thermal tolerance have been studied intensively, and in general, heat shock proteins are involved in tolerance responses in many organisms (Feder and Hofmann, 1999). In fungi thermal tolerance is mediated by heat shock proteins (Piper, 1993; Glatz *et al*., 2015), other metabolites in particular the sugar trehalose (De Virgilio *et al*., 1994; Bonini *et al*., 1995; Argüelles, 1997), and changes in lipid bilayer composition (Glatz *et al*., 2015). However, the regulation thermotolerance is not completely understood. Epigenetic mechanisms are one possibility for indirectly mediating the influence of past environments. These include chromatin modifications such as DNA methylation (Bird, 2002), and different types of histone modifications (Bannister and Kouzarides, 2011). It is known that epigenetic mechanisms can mediate cell memory (Bird, 2002; Gaydos *et al*., 2014; de la Paz Sanchez *et al*., 2015), plasticity (Kooke *et al*., 2015; Kronholm *et al*., 2016), and are required for certain parental effects (Wibowo *et al*., 2016) or between generation plasticity (Luna and Ton, 2012; Rasmann *et al*., 2012). Effects that are passed from one generation to the next can create lag in fluctuating environments. However, memory effects can also occur within one generation. If epigenetic mechanisms control phenotypic plasticity, they may be needed in fluctuating environments. In particular, we have previously observed that the *set-2* mutant, which is deficient in histone 3 lysine 36 methylation, and the *qde-2* mutant, which is defective in milRNA processing, are both impaired in their temperature responses in constant temperatures (Kronholm *et al*., 2016).

This study has two aims: the first is to investigate if reaction norms measured in constant temperatures can be used to explain the growth rate of the filamentous fungus *Neurospora crassa* in fluctuating temperatures, and if not, why do reaction norms measured in constant environments fail? The second is to investigate the role of epigenetic mechanisms within a generation in fluctuating environments: does the absence of certain chromatin modifications change the shape of reaction norms in fluctuating environments, indicating that this specific mechanism is needed in a fluctuating environment? In order to investigate the importance of epigenetic mechanisms in response to fluctuating temperatures, we used a collection of deletion mutants that each lacked a certain epigenetic mechanism and grew these mutants in the different fluctuating environments. The hypothesis is that if an epigenetic mechanism is involved in sensing temperature change or in the expression of acclimation, mutants deficient in this mechanism would acclimate slower or ineffectively. If an epigenetic mechanism is involved in memory of past environments and causes lag in acclimation to a new environment, then mutants deficient in this mechanism would acclimate faster.

As a fluctuating environment, we used two temperatures: 30 and 40 °C and durations ranging from 30 to 720 minutes the cultures spent in each temperature. The species *N. crassa* occurs mainly on burned vegetation in tropical and sub-tropical regions (Turner *et al*., 2001), and its optimum growth temperature is around 35 °C, while 40 °C is a stressful temperature for it (Kronholm *et al*., 2016). We also investigated how long it takes for the fungus to acclimate from one temperature to another, that is, to reach a constant growth rate in the new temperature. We then tested different growth models to predict growth in fluctuating environments, such as models with no lag in acclimation or lag effects included.

We find that reaction norms measured in constant environments predict growth rate poorly in rapidly fluctuating environments, due to time lag in temperature acclimation. Better predictions are obtained from a model that takes lag times in to account. Epigenetic mechanisms, in particular histone modifications H3K36 methylation and H3K4 trimethylation, affect growth in fluctuating environments, but are not as important as in constant environments. We also found that H3K4 trimethylation and RNA interference pathway are involved in acclimation responses.

## Materials and methods

### Neurospora strains and growth measurements

We previously obtained a set of deletion mutants from the *Neurospora* knockout mutant collection (Colot *et al*., 2006) from Fungal Genetics Stock Center (FGSC) (McCluskey *et al*., 2010), the *qde-2* mutant was provided by Tereza Ormsby and Eric Selker, and backcrossed these mutants to the wild type strain 2489 (Kronholm *et al*., 2016). Back-crossing was done so that mutants differed from the control genotype only in the deleted gene. Mutants used in this study are listed in table 1, all strains were of mating type A. Procedures used for backcrossing and confirming the presence of the deletions by PCR can be found in Kronholm *et al*. (2016). The experiment contained 20 mutants and the wild type strain 2489, 21 strains in total.

**Table 1:**
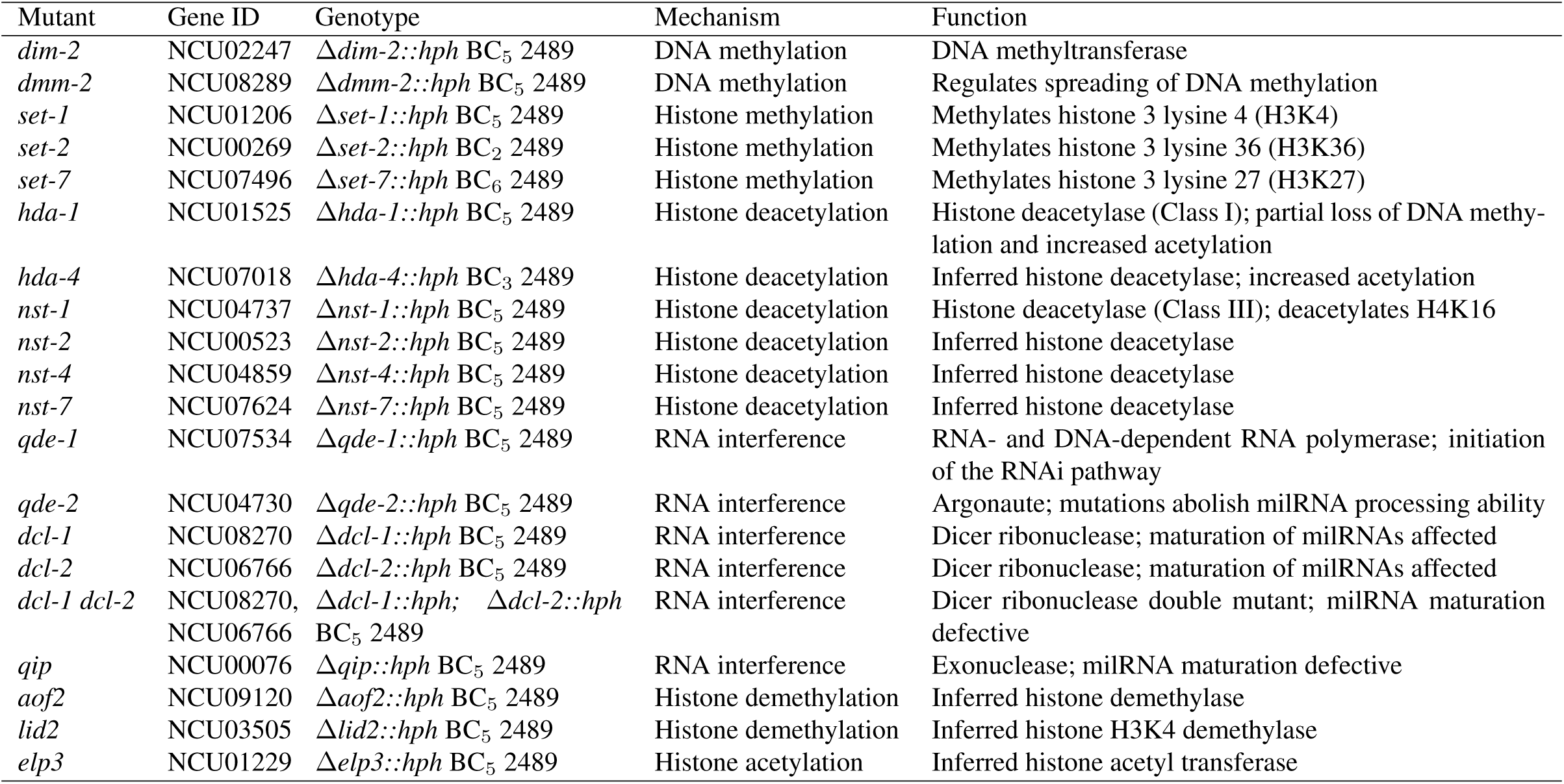
Gene ID is based on genome assembly NC12. The function column describes the biochemical activity of the protein. milRNA = microRNA-like RNA. BC = backcross. 2489 is the control genotype and the other parent for the backcrosses.

We grew the strains on Vogel’s standard growth medium (Metzenberg, 2003) with 1.5% agar in disposable 25 ml pipettes prepared by the method of White and Woodward (1995). At the start of each experiment a strain was inoculated at the other end of the pipette and the growth of a strain was followed by marking the position of the mycelial front at a given time point. Growth rate was obtained as the slope of the regression line for time againts distance the mycelium had grown. Detailed description of the growth measurements can be found in Kronholm *et al*. (2016).

### Growth in fluctuating environments

To measure growth rate in fluctuating environments, we used programmable growth chambers (MTM-313 Plant Growth Chamber, HiPoint Corp., Taiwan) to change the temperature periodically. We used fluctuations with an amplitude of 10 °C from 30 °C to 40 °C. We used eight different fluctuation regimes: cultures spent either 30, 60, 90, 120, 150, 240,480, or 720 min in one temperature during one cycle. Thus fluctuations had periods of 1, 2, 3,4, 5, 8, 16, or 24 h. Throughout this study, when discussing the different fluctuation regimes, we will refer to the time cultures spent in one temperature as duration of a step. Growth was followed for a period of 104 h and measurements were taken every 8 and 16 hours to obtain growth rates for different step durations (Supplementary figure S1A). In each fluctuation regime temperature changed in a stepwise manner (Supplementary figure S2), so that during one cycle cultures spent half of the time in 30 ^°^C and half of time in 40 ^°^C. Thus the mean temperature for each treatment was 35 ^°^C and the total time spent in 30 and 40 ^°^C the same.

We grew the 21 strains in the 8 different fluctuation regimes with 5 replicates each, the strain *nst-7* was missing from step durations 240, 480, and 720 min, giving 825 growth assays in total. With two growth chambers with three independently controlled compartments each, we had six different compartments available. To control for compartment effects, we rotated the fluctuation regimes among the compartments between different replicates. Strains were always randomised within a compartment.

In a preliminary experiment we measured the rate of temperature equilibration between growth chamber air and the agar tubes used to measure growth rates of the strains. We inserted a temperature probe into the agar of a growth tube and monitored the temperature in the growth tube and the air temperature in the growth chamber during heating and cooling. It took about twelve minutes for the growth chamber air temperature to rise from 30 to nearly 40 °C, while the same temperature change took approximately 20 min for the agar (Supplementary figure S3). During heating the growth chamber first raised the temperature nearly to the set point, but to prevent overshooting, raising the temperature the last remaining degree took most of the elapsed time. Cooling was more efficient and it took approximately six minutes for the agar temperature to drop from 40 to 30 ^°^C. Therefore the actual times the cultures spent in 40 and 30 ^°^C were shorter than the nominal temperature settings, and cycles of 30 min at each temperature were the shortest fluctuations possible that could be achieved with our system.

### Temperature shift experiments

We performed temperature shift experiments to measure how the growth rate changed, and if there was a lag time, when *Neurospora* acclimated to a new temperature. In a shift experiment we initially inoculated cultures and grew them in either constant 30 or 40 °C for 24 h to allow the cultures to acclimate to the current temperature. Then we swapped the cultures between the two different incubators, those cultures that had been initially growing in 30 ^°^C were moved to 40 ^°^C and vice versa. As the growth chambers were constantly at the same temperature and the cultures were shifted, there was no time lag associated with heating or cooling the growth chamber, only the heat transfer to agar medium itself. After the shift we recorded the growth of the cultures every hour for 8 hours and for two additional time points, 25 and 32 h, to allow fine grain monitoring of changes in growth rate after the shift (Supplementary figure S1B). For analysis we calculated growth rates for two hour intervals as recording one hour of growth in the tubes was difficult as the marks were very close to each other and thus the data measured every two hours provided a reasonable smoothing of measurement noise.

We performed two shift experiments. In the first experiment we used two genotypes: the wild type control genotype 2489 and the *qde-2* mutant, both genotypes were replicated 10 times in both shifts giving 40 growth assays in total. We included *qde-2* in this initial experiment as our previous results suggested that it is involved in temperature responses (Kronholm *et al*., 2016). In the second expereriment we performed a temperature shift experiment with all of the mutants except *nst-7, dcl-1*, and *dcl-2*. The *dcl* single mutants were excluded because in all previous experiments they had the same phenotype as the *dcl-1 dcl-2* double mutant. In the second experiment each of the 18 genotypes was replicated 4 times in both shifts giving 144 growth assays in total.

### Estimating the cell cycle durations

*Neurospora* is a filamentous organism, its cells remain cytoplasmically connected to each other and the nuclei can move within the mycelium. Thus, asexual mitotic divisions do not constitute generations as they do in unicellular microbes and estimating the number of mitotic divisions that occurred for a certain amount of growth is not simple. In addition, mitotic divisions are not fully synchronous, and it is not known how many of the nuclei are actively dividing in a growing hyphae. The duration of cell cycle has been estimated to be 103 min (Martegani *et al*., 1981) in conditions that correspond to a growth rate of 4.4 mm/h (Ryan *et al*., 1943; Kronholm *et al*., 2016), and around 217 min (Martegani *et al*., 1981) in conditions where growth rate is around 2.3 mm/h (Ryan *et al*., 1943). Based on these numbers we interpolated cell cycle durations for our observed growth rates assuming a linear relationship. These numbers should be regarded as rough estimates, that are only meant to give an approximate idea of what is the relationship between step duration and cell cycle duration.

### Data analysis

#### Analysis of growth rates in fluctuating environments

We used a linear mixed model to investigate the effects of the different fluctuation regimes on the growth of the different strains. We first fitted the growth chamber compartment effect to the data and removed the compartment effect by subtracting the effect from the raw data. Because the strains and treatments were randomly distributed, removing the average effect of compartment does not remove strain or treatment effects. We encoded the different step durations as factors to allow us to fit the same model for all strains, as responses were non-linear. The model was

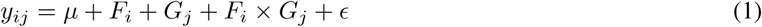

where *µ* is the intercept, *F_i_* is the *i*th step duration, *G_j_* is the *j*th genotype, *∊* is the residual, and *y_ij_* is the growth rate of the *j*th genotype in step duration *i*. The duration of the step was fitted as a fixed effect and genotype as a random effect. The mixed model was fitted with the “lmer” function in R (R Core Team, 2013) and statistical testing performed with the “lmerTest” package (Kuznetsova *et al*., 2015). The lmerTest package implements *F*-tests using type III sums of squares with Satterthwaite correction for degrees of freedom. For comparing reaction norm shape for each mutant to the control, we used a pairwise ANOVA to test if the genotype by step duration interaction was significant. Correction for multiple testing was done using the Bonferroni-Holm method (Holm, 1979).

#### Estimating lag times from temperature shift experiments

To obtain lag times from the temperature shift experiments we fitted non-linear regression curves to the temperature shift data, for each direction of temperature change, using the model

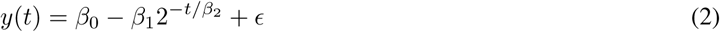

where *y* is the growth rate, *β*_0_ is the asymptote or the final growth rate after acclimation, *β*_1_ is the amount of growth rate to be gained (i.e. *β*_0_ − *β*_1_ is the growth rate immediately after temperature change), *β*_2_ is the time it takes for growth rate to increase to half of its maximum value, *t* is the time elapsed since temperature change, and *∊* is the residual (Venables and Ripley, 2002). This yielded a set of parameters *β*_0_, *β*_1_, and *β*_2_ estimated for each direction of temperature change. Fitting a negative exponential function has the advantage that all the parameters have a biological interpretation. We are interested in the *β*_2_ parameter as it gives us a measure for the length of the lag time. When fitting the model to the shift from 30 to 40 ◦C data we excluded data points from constant 30 ◦C and assumed that growth rate drops faster than the resolution in our data collection. For fitting the model and testing for differences in lag times among the different genotypes we used the “nls” and “gnls” functions in R.

#### Growth modeling based on temperature shift experiments

To model growth in fluctuating environments for the control genotype, 2489, we used the acclimation times obtained from temperature shift experiments, and tested if could predict growth in fluctuating environments with this data. After obtaining numerical estimates for equation 2 for both acclimation directions, we defined two lag functions giving the instantaneous growth rate as a function of time *t* elapsed since temperature change as

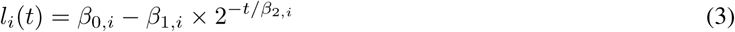

where *i* = 1 for a change from 30 to 40 °C and *i* = 2 for a change from 40 to 30 °C. Then we used growth rate models to predict growth in fluctuating environments. In our initial model, growth rate *r_p_* in a fluctuating environment where step duration was *p* hours during one cycle was

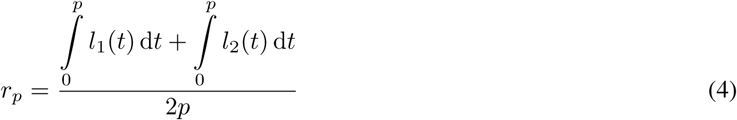

where the numerator gives the amount of growth (in mm) that occurs during one temperature cycle, *l*_1_(*t*) and *l*_2_(*t*) are the lag functions obtained from fitting equation 2 to the different temperature shifts and using the obtained *β* parameters. We integrate over the time spent in given temperature to obtain the predicted growth that occurs after temperature shift, the denominator gives the total time (in h) for one full temperature cycle to obtain the growth rate as mm/h.

As our initial lag model worked poorly for fast fluctuations (see results) we later refined our inital model to account for partial acclimation. The predicted lag times were much longer than the step durations in fast fluctuations. Therefore, acclimation happens only partially before the temperature changes again and consequently next acclimation does not begin from the growth rate the culture would eventually reach in constant temperature. We modified the model so that we updated the parameter *β*_1_, setting the amount of growth rate to be gained in the next cycle to reflect the growth rate reached in the previous temperature.

The refined model followed the following algorithm: a culture spent p hours first in 30 °C and we calculated the distance grown, then we calculated the current growth rate, *r*\*, at time *p*. If *r*\* *< β_0_* of the next lag function (*β*_0_,_2_ during first cycle), then we set 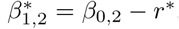, otherwise we set 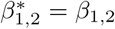. We then calculated the distance grown using the updated *l*_2_ function. After *p* hours of growth in the next temperature, current growth rate was again calculated and it was checked if *r*\* < *β*_0,1_ then we set 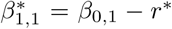, and to 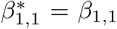 otherwise. We then calculated the distance grown using the updated *l*_1_ function. The empirical measurements for the different strains in the fluctuating environments were done over 103 hours, so the number of temperature cycles in the model, *n*, was determined by how many full cycles of 2*p* could fit into 103 hours. The growth rate was obtained from

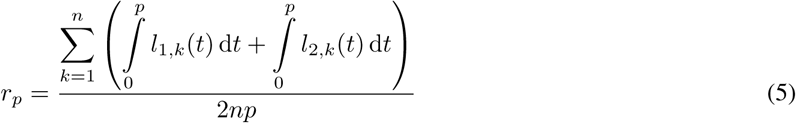

where the *β*_1_*_,i_* parameters were updated for each step *k* as described above.

We compared how well the simulation models fit the observed data by calculating mean squared deviations from the observed data for each model. We can use mean squared deviations (MSD) for comparing the relative ranks of the different models, but MSD does not tell us how well the model fits the data in an absolute sense. We chose to use MSD because traditional measures of model fit, like *R*^2^ or AIC, cannot be calculated for our simulated models.

## Results

### Effect of temperature fluctuations on growth

At constant 30 °C the control genotype 2489 grows at 4.4 mm/h and at constant 40 °C at 2.4 mm/h (Kronholm *et al*., 2016). The control strain has its optimum at 35 °C (Figure 1A). We observed that the growth rate of the control strain in step durations from 30 to 150 min was mainly determined by its growth rate in constant 40 °C (Figure 1B), as growth rate of 2489 in these step durations was only slightly above 2.4 mm/h. Growth rate increased slightly with the period of fluctuations. Under the assumption that acclimation to 30 and 40 °C would be instantaneous and growth rate would change immediately to corresponding growth rate at constant temperature, the growth rate per hour in fluctuating temperature would be average of the growth rates in constant 30 and 40 ^°^C because under all fluctuation regimes the cultures spend equal amount of time in 30 and 40 ^°^C. This naive expected growth rate would be 3.38 mm/h, the observed growth rates in these fast fluctuations were well below this rate (Figure 1B). When step durations increased to 240 min and above growth rate increased more rapidly, and at step durations 480 and 720 min growth rate was very close to the expected growth rate of 3.38 mm/h.

**Figure 1:**
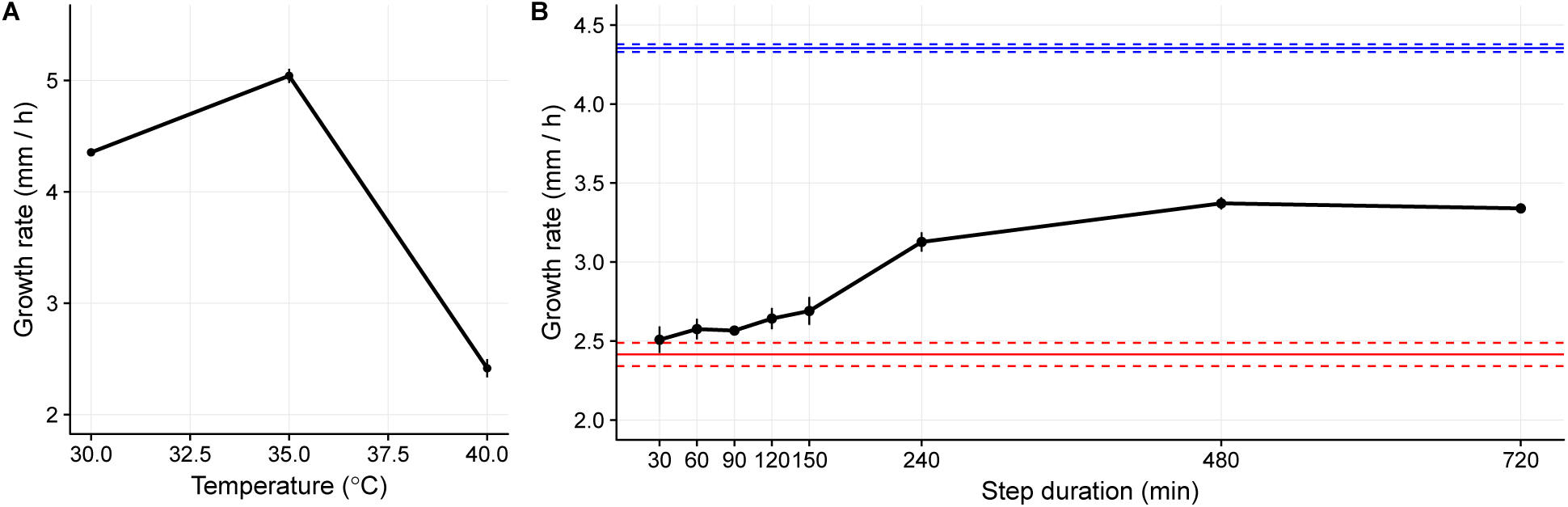
A) Reaction norm for the genotype 2489 in constant temperatures, n = 12 for each point. Data from Kronholm et al. (2016). B) Reaction norm for the genotype 2489 in different step durations when temperature fluctuates between 30 and 40 °C, n = 5 for each point. Horizontal blue and red lines show growth rate at constant 30 and 40 °C, respectively. Error bars and dashed lines show means ± SE.

The estimated average cell cycle durations were 206, 202, 202, 200, and 195 min for step durations of 30, 60, 90, 120, and 150 min respectively. With these step durations each step durations was shorter than the lenght of the complete cell cycle. The estimated average cell cycle durations were 172, 157, and 162 min for step durations of 240, 480, and 720 min. For these longer step durations multiple cell cycles happened within one step.

### Effects of epigenetic mechanisms in fluctuating environment

Next we tested if epigenetic mechanisms were important for growth in fluctuating environments. Growth rates of most strains were around 2.51 mm/h with step duration of 30 min and increased to 2.70 mm/h when step duration increased to 150 min. With longer step durations growth rate further increased to around 3.34 mm/h (Figure 2). In a mixed model the effect of step duration was significant (F = 383.14, numerator df = 7, denumerator df = 137.7, p < 2.2 × 10^−16^), showing that different step durations affected the growth of the strains in fluctuating environments. Reaction norms for growth in fluctuating environments of the genotypes show that in general there was a small increase in growth rate for short step durations until the 150 minute step, and then growth rate increased faster as the strains had more time to acclimate in the 240 and 480 minute steps but growth rate plateaued after that (Figure 2). We also tested whether the small increase in growth rate observed for the short step durations was significant by analysing the steps from 30 to 150 minutes separately. In this model the effect of duration of the step was significant (F = 18.69, numerator df = 4, denumerator df = 489.01, p = 2.62 × 10^−14^), indicating that while the effect of duration of the step in short timescales may be small, it was likely a real effect.

**Figure 2:**
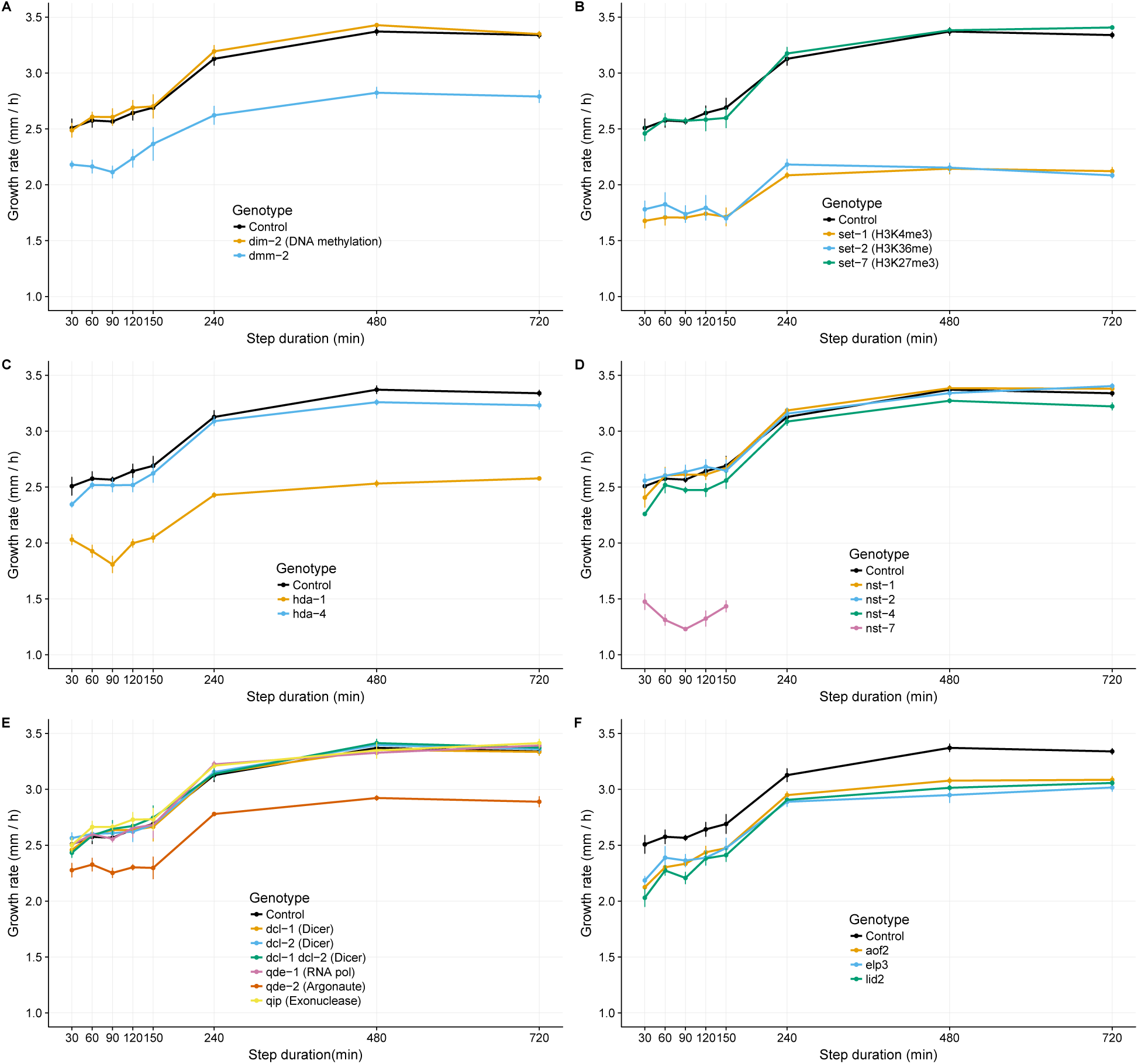
Reaction norms of different classes of mutants. Lines and error bars show means ± SE, n = 5. A) DNA methylation mutants. B) Histone methylation mutants. C) Histone deacetylation class I mutants. D) Histone deacetylation class III mutants. E) RNA interference mutants. F) Histone demethylation and acetylation mutants.

There were differences in reaction norm elevation among the different genotypes, as shown by a significant genotype effect (χ^2^ = 387.62, df = 1, p = 2.73 × 10^−86^). Differences in reaction norm elevation indicate that some mutants grew slower overall (Figure 2), some of the mutants were undoubtedly impaired in their normal cellular metabolism and grew slower as a consequence. However, we were interested in differences in reaction norm shape, which indicated that responses to the environment were different in the different strains. There was evidence that reaction norm shapes were different as indicated by a significant step duration × genotype interaction (χ^2^ = 17.18, df = 1, p = 3.4 × 10^−5^). Some strains responded to the environment differently, suggesting that epigenetic mechanisms affect growth rate in fluctuating temperatures. To identify which mutants reacted differently than the control strain, we performed a pairwise ANOVA where we tested each mutant separately (Table 2).

**Table 2:** Comparing all mutants to the control in a pairwise ANOVA. Differences in reaction norm elevation from the control are tested with the genotype effect, differences in reaction norm shape are tested with the genotype step duration interaction effect (G × E). For genotype effect df = 1, for step duration df = 7, for G × E df = 7, and for residuals df = 59. Significant G × E term indicates change in reaction norm shape, p-adj. indicates a p-value adjusted for multiple testing using the Bonferroni-Holm method.

| Genotype | F-value<br>Genotype | p-value<br>Genotype | p-adj.<br>Genotype | F-value<br>$G \times E$ | p-value<br>$G \times E$ | p-adj.<br>$G \times E$ |
| --- | --- | --- | --- | --- | --- | --- |
| *aof2* | 61.073 | 1.13E-10 | 1.46E-09 | 0.654 | 0.709 | 1 |
| *dcl-1* | 0.000 | 0.997 | 1 | 0.163 | 0.992 | 1 |
| *dcl-2* | 0.582 | 0.448 | 1 | 0.273 | 0.962 | 1 |
| *dcl-1 dcl-2* | 0.482 | 0.49 | 1 | 0.081 | 0.999 | 1 |
| *dim-2* | 0.958 | 0.331 | 1 | 0.108 | 0.998 | 1 |
| *dmm-2* | 155.624 | 8.58E-19 | 1.37E-17 | 0.803 | 0.588 | 1 |
| *elp3* | 56.694 | 3.43E-10 | 4.11E-09 | 0.805 | 0.587 | 1 |
| *hda-1* | 641.911 | 4.52E-35 | 7.68E-34 | 2.025 | 0.0654 | 1 |
| *hda-4* | 10.298 | 0.00208 | 0.0208 | 0.303 | 0.95 | 1 |
| *lid2* | 128.872 | 5.63E-17 | 7.88E-16 | 1.021 | 0.426 | 1 |
| *nst-1* | 0.020 | 0.887 | 1 | 0.389 | 0.906 | 1 |
| *nst-2* | 1.181 | 0.281 | 1 | 0.223 | 0.978 | 1 |
| *nst-4* | 19.003 | 4.83E-05 | 0.000532 | 0.722 | 0.653 | 1 |
| *nst-7* | 1960.351 | 5.85E-43 | 1.17E-41 | 2.094 | 0.0948 | 1 |
| *qde-1* | 0.242 | 0.624 | 1 | 0.252 | 0.97 | 1 |
| *qde-2* | 144.391 | 4.65E-18 | 6.98E-17 | 1.019 | 0.427 | 1 |
| *qip* | 3.320 | 0.0731 | 0.658 | 0.289 | 0.956 | 1 |
| *set-1* | 1163.202 | 9.14E-43 | 1.74E-41 | 3.728 | 0.00191 | 0.0364 |
| *set-2* | 783.529 | 1.29E-37 | 2.32E-36 | 4.396 | 0.000491 | 0.00981 |
| *set-7* | 0.053 | 0.818 | 1 | 0.393 | 0.903 | 1 |

### Estimating acclimation times

Growth rate in short term fluctuations could not be explained by the naive model of instant acclimation, and was close to growth rate in constant 40 °C. Thus, we suspected that there was a lag time in acclimation response. To estimate how fast growth rates changed we performed temperature shift experiments, where we shifted cultures from 30 to 40 ^°^C and vice versa and monitored how growth rate changed. In the first experiment, when cultures were shifted from 30 to 40 °C growth rate initially dropped to around 2 mm/h and then recovered to around 2.6 mm/h (Figure 3A). This took around 6 hours, there was an increase in the growth rate around 8 hours, but later the growth rate seemed to stabilise to slightly lower value. The origin of this transient increase is unknown, it showed up earlier also in the *qde-2* mutant, so it probably cannot be attributed to a random fluctuation in the growth chamber conditions. However, the overall pattern of low initial growth rate after the switch and subsequent recovery was clear. Lag time parameter, 0_2_ ± SE, obtained from a non-linear fit was 1.38 ± 0.34 h for genotype 2489, and 0.48 ± 0.36 h for *qde-2*. This difference of 0.9 h was suggestive but not significant (t = −1.84, df = 113, p = 0.0691). When cultures were shifted from 40 to 30 °C growth rate recovered to the growth rate at constant 30 °C but this took up to 8 hours (Figure 3B). Lag time parameter, *β*_2_ ± SE, obtained via non-linear fit was 2.06 ± 0.18 h for control genotype 2489, and 3.56 ± 0.27 h for genotype *qde-2*. The difference of 1.49 h was significant (t = 4.57, df = 130, p = 1.11 × 10^−5^). These data show that temperature acclimation in *Neurospora* is not instantaneous but takes up to 6 to 8 hours. Furthermore, acclimation is asymmetric; the direction of environmental change matters, as recovery from high temperature took longer than acclimation to high temperature. Acclimation times were much longer than time it took for growth chambers to equilibrate to a certain temperature (Supplementary figure S3), so temperature transfer cannot explain asymmetries in acclimation. Control and *qde-2* differed in their responses, in particular the *qde-2* mutant recovers slower from high temperature than the control strain (Figure 3B). Acclimation times were also longer than estimated cell cycle times of 1.7 and 3.6 h for 30 or 40 ^°^C respectively. Thus, acclimation takes approximately from 2 to 5 cell cycles.

**Figure 3:**
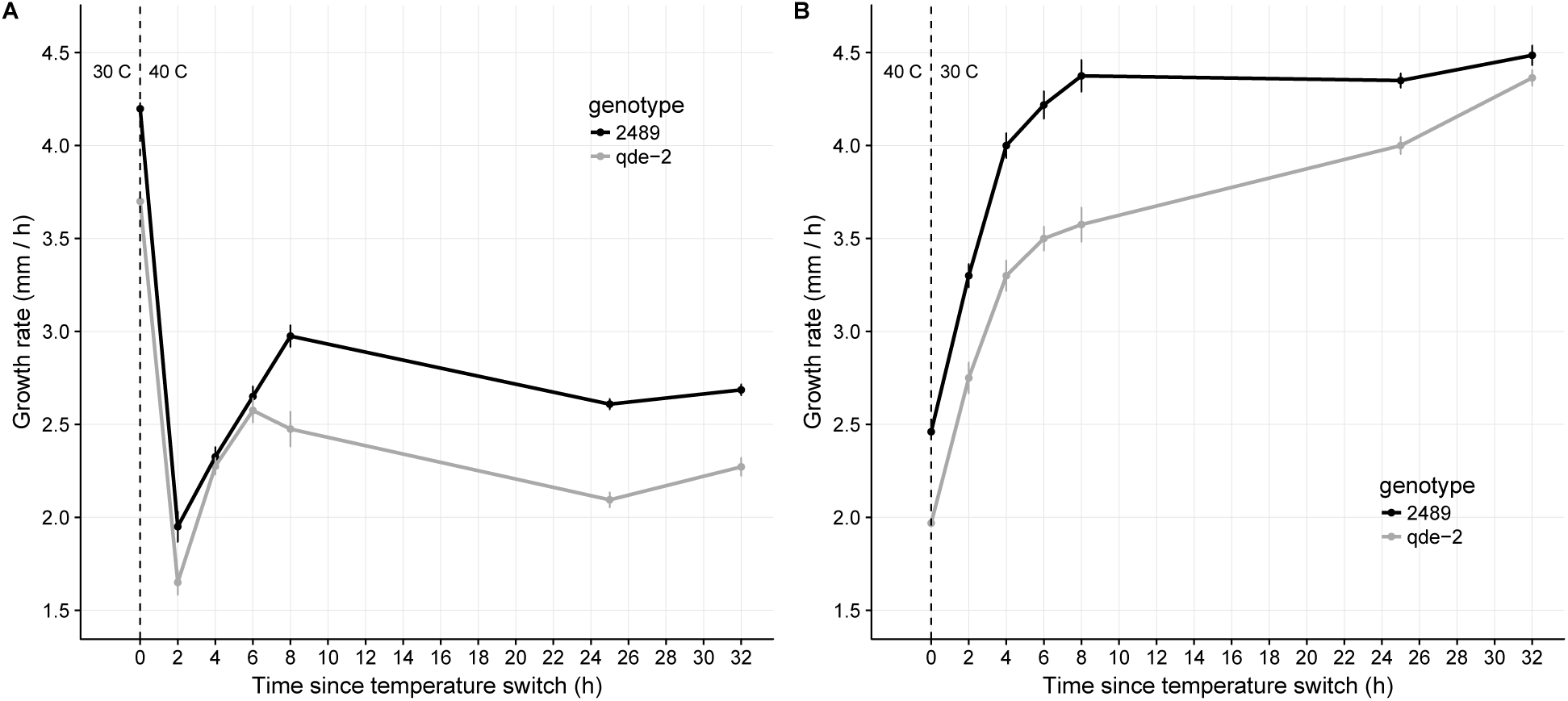
Changes in growth rate after temperature shift. Lines and error bars show means ± SE, n = 10. A) Temperature shift from 30 to 40 °C. B) Temperature shift from 40 to 30 °C.

We then performed a second temperature shift experiment, where we estimated lag times for all of the mutants. Estimates of lag parameter *β*_2_ were similar in shift from 30 to 40 °C to first experiment for the control genotype. None of the mutants had significantly different lag parameters from the control, even though some, such as *hda-1* had increased variation (Figure 4). For the *set-1* mutant the lag parameter could not be estimated as the model could not be fitted, as its growth rate did not recover from the intial depression after the transfer (Figure S4). For the shift from 40 to 30 °C the lags were somewhat lower than in the first lag experiment, the lag parameter *β*_2_ was 1.79 for the control. The *set-1* mutant had a longer lag *β*_2_ = 3.73 ± 0.35 and difference to the control was significant (t = 3.57, df = 443, p = 4.02 × 10^−4^. In this experiment, the *qde-2* mutant also had a longer lag time than the control (Figure 4) but this difference was not significant (t = 1.62, df = 443, p = 0.1067). It is likely that this is because the second experiment had less statistical power. For the other mutants lag times were not different from the control (Figures 4 and S5). Next we will look at growth in fluctuating environmen and acclimation responses of the different mutant classes in more detail.

**Figure 4:**
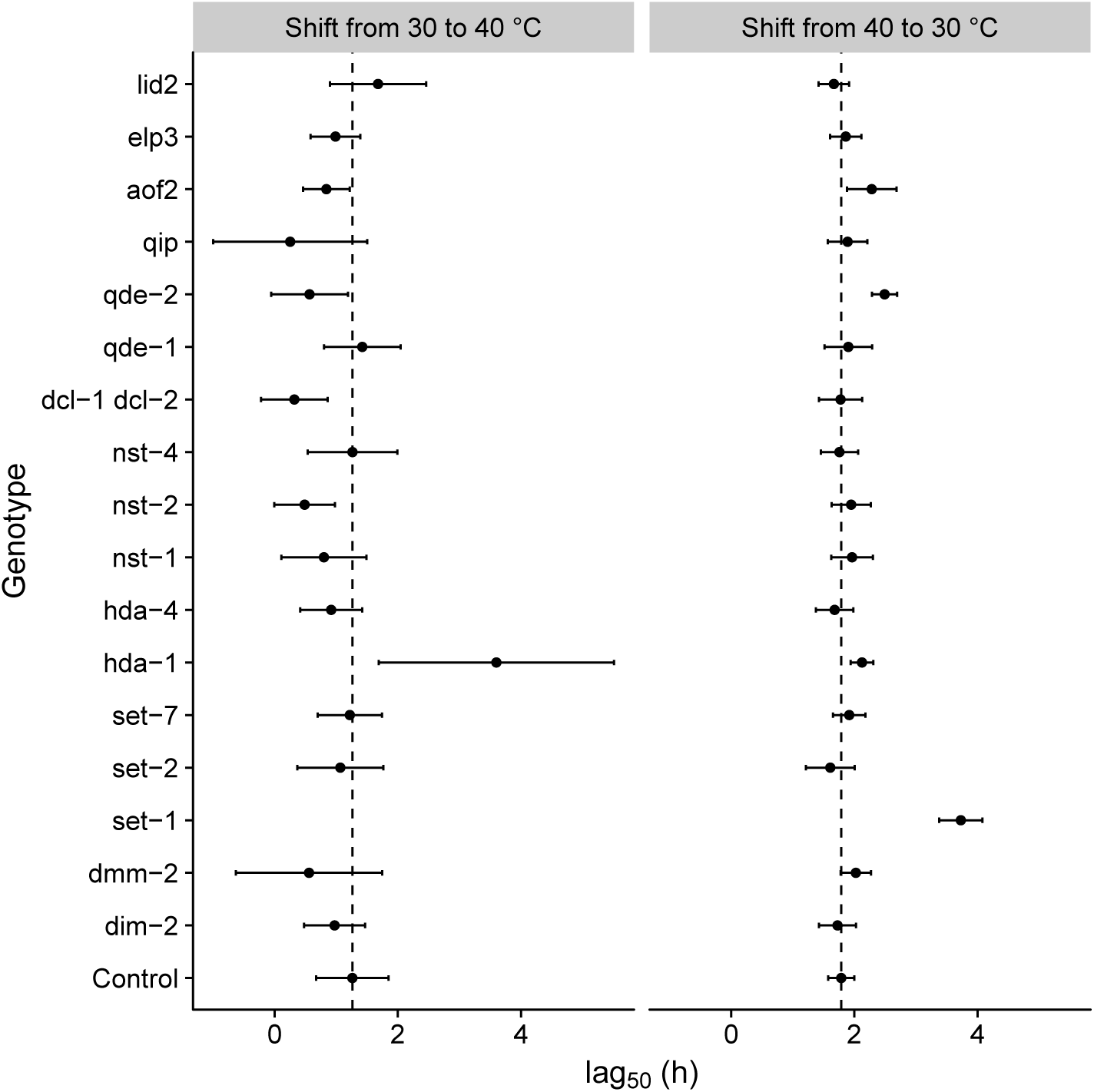
Lag time parameter *β*_2_ values for all mutants. Points show an estimate from a non-linear model ± SE, n = 4. Dashed vertical line shows the estimate of the control genotype 2489.

#### DNA methylation

Reaction norms for growth rate in fluctuating environments of strains with deficiencies in DNA methylation are shown in figure 2A. The *dim-2* mutant lacks a DNA methyltransferase enzyme but showed no difference to the control. Indicating that DNA methylation is not important for temperature response in *Neurospora*. The *dmm-2* mutant is deficient in containing DNA methylation to heterochromatic regions, and as a consequence DNA methylation spreads to normally unmethylated regions in this mutant. It grew slower than the control but responded to different step durations the same way as the control. Neither mutant had any differences in acclimation (Figures 4, S4A, S5A). This is consistent with our previous observation that neither mutant had a any differences compared to control in their temperature response measured in constant temperatures (Kronholm *et al*., 2016).

#### Histone methylation

We tested three different mutants with deficiencies in histone methylation, *set-1* which lacks H3K4 trimethylation, *set-2* which is impaired in H3K36 methylation, and *set-7* which lacks H3K27 trimethylation. The growth of *set-7* mutant was not different from the control (Figure 2B, 4), but both *set-1* and *set-2* had lowered reaction norm elevation. Furthermore both *set-1* and *set-2* differed from the control in reaction norm shape (Table 2). Both genotypes had reached their maximal growth rate already at the 240 min step, and during the short fluctuations they did not increase their growth rate as much as the control (Figure 2B). We had previously observed that the temperature response of *set-2* is altered (Kronholm *et al*., 2016), which may explain its response in fluctuating environments. For *set-2* there were no significant differences in acclimation (Figure 4). We evaluated if changes in reaction norm shape could be the result of this strain reaching its steady-state growth rate faster if it has to acclimate less than the control, even if acclimation happens at the same rate. We calculated the expected growth rate under the assumption of no lag from their growth rates at constant temperature, data obtained from Kronholm *et al*. (2016). For *set-2* growth rate in 720 min step was 2.08 mm/h and this was exactly at the no lag expectation of 2.08 mm/h. Thus, for *set-2* it is possible that differences in reaction norm shape reflect just lower acclimation requirement. Acclimation for *set-1* took longer in the 40 to 30 °C switch (Figure 4). However, for the 30 to 40 °C switch the model could not be fitted as it appears there is almost no acclimation happening for *set-1* (Figure S4B). This could indicate that *set-1* is required for acclimation to 40 ^°^C.

#### Histone deacetylation

We tested two mutants of the histone deacetylase class I genes: *hda-1* and *hda-4*, and four mutants of the histone deacetylase class III genes: *nst-1, nst-2, nst-4*, and *nst-7*. The *hda* mutants grew slower than the control (Figure 2C, Table 2), but there were no significant differences in reaction norm shape or acclimation time (Figure 4). The *hda-1* had higher acclimation time and higher variability in 30 to 40 °C acclimation, but this effect was not significant. This is probably a reflection of poor model fit, as the shape of the acclimation shows very little acclimation (Figure S4C). So it remains possible that *hda-1* has some effect. For the *nst* mutants, *nst-1* and *nst-2* did not differ from the control, *nst-4* had slightly lower elevation than the control but same shape, and *nst-7* grew very poorly overall indicating severe problems in normal cellular functioning (Figure 2D, Table 2). No differences in acclimation were observed for the *nst* mutants (Figure 4, S4D, S5D).

#### RNA interference

*Neurospora* produces a diverse set of different small RNAs, we tested six mutants in the RNA interference pathway; including the two Dicer ribonuclease genes *dcl-1, dcl-2* and their double mutant *dcl-1 dcl-2, qde-1, qde-2*, and *qip*. The growth of all other mutants except *qde-2* did not differ from the control (Figure 2E). The *qde-2* mutant grew overall slower than the control, but its reaction norm shape was not different from the control (Table 2). In short step durations, it looked like *qde-2* did not increase its growth rate like the control, but this effect was not significant. In the temperature shift experiments *qde-2* acclimated slower than the control in the 40 to 30 °C shift (Figure 3, 4, S5E), which may explain the reaction norm trend even if there was a lack of power.

#### Histone demethylation and acetylation

The ELP3 protein is an inferred histone acetyl transferase, while LID2 and AOF2 are inferred histone demethylases. All three mutant strains had a lower growth rate in fluctuating environments than the control (Figure 2F), but they had the same reaction norm shape as the control (Table 2). This was expected based on our previous results, as we found no differences in the temperature response of these mutants (Kronholm *et al*., 2016). Furthermore, no differences in acclimation were observed for these mutants in constant temperatures (Figure 4, S4F, S5F).

### Modeling growth in fluctuating environments

The long lag times suggest a mechanistic explanation for slow growth rate in rapidly fluctuating environments. We tested this by fitting a growth model based on the observed lag times. We first fitted non-linear regressions to the lag data to estimate empirical parameters for the lag functions (Equation 3, Table 3). Then we used those lag functions to predict the growth rates in fluctuating environments with different step durations. For the initial lag model we calculated predicted growth rates by integrating over the times the culture spent in the different temperatures. Comparing the predictions from the initial lag model to the entire observed data set showed that the initial lag model worked poorly (Figure 5), particularly in short step durations. The initial model probably failed because in short step durations there was not enough time to acclimate completely and reach the expected growth rate. We then refined the initial model to account for partial acclimation during the short time intervals, and the refined model fitted the data much better (Figure 5). To assess how well each model fitted the data, we calculated mean squared deviations for each predicted value from the observed data as a measure of model fit for the two different lag models, naive expectation of no lag, and for growth rate in constant 40 °C. The MSD values were: 0.4 for the model with no lag, 0.31 for the initial lag model, 0.31 for the growth rate in constant 40 °C, and 0.02 for the refined model. All the other models were clearly inferior compared to the refined lag model. These results indicate that growth rate of *Neurospora* in the fluctuating environments can be predicted to a reasonable degree if the lag functions are known.

**Table 3:** Empirical estimates for lag function parameters used in the models. Obtained by fitting equation 2 to the temperature shift data.

| Temperature change | Notation | $\beta_0$ | $\beta_1$ | $\beta_2$ |
| --- | --- | --- | --- | --- |
| 40 to 30 °C | $l_1$ | 4.45 | 2.03 | 2.06 |
| 30 to 40 °C | $l_2$ | 2.74 | 2.25 | 1.38 |

**Figure 5:**
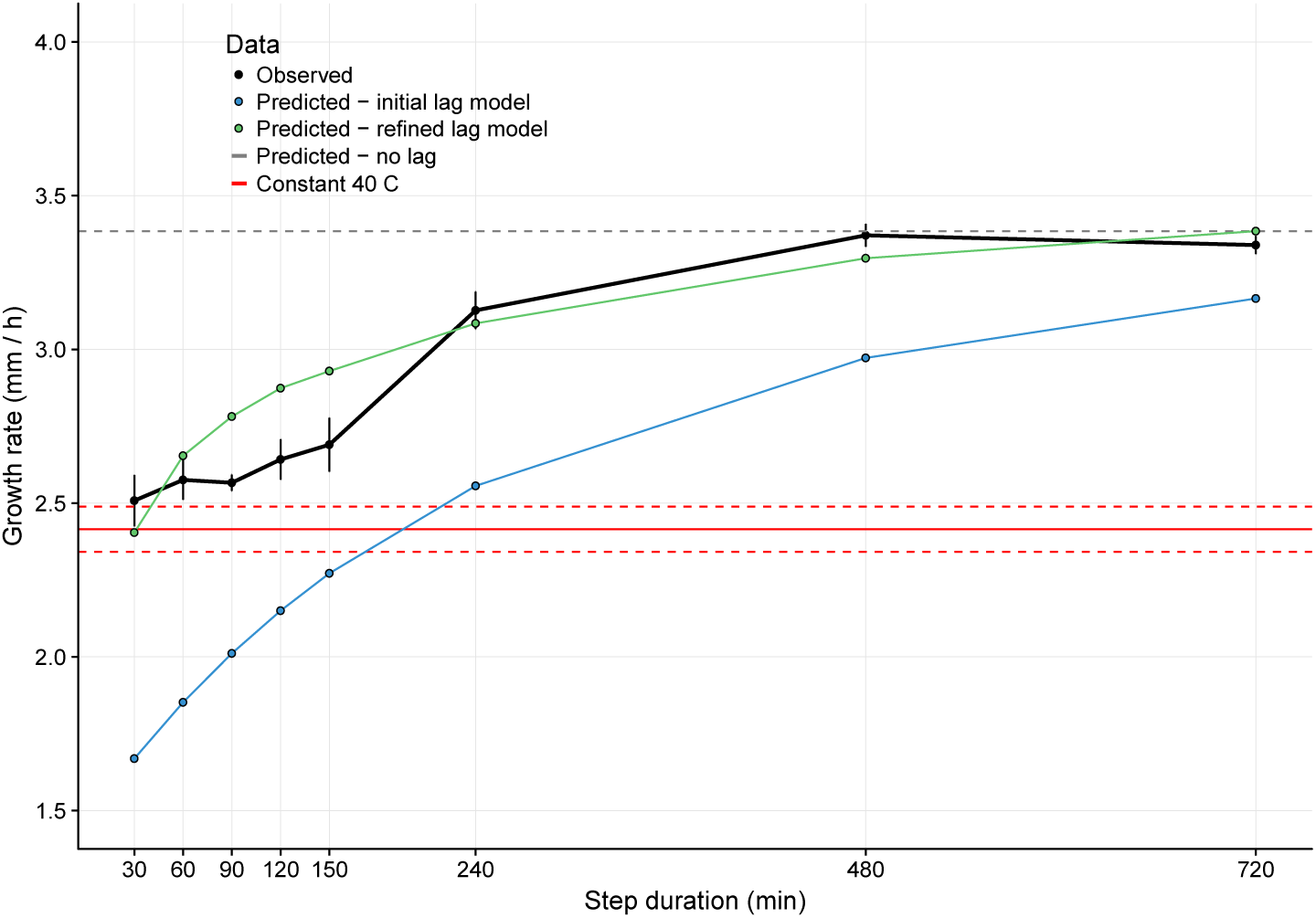
Comparison of how well different growth models with lag fit to the observed fluctuation data of genotype 2489. For the observed data, points are means ± SE, n = 5. Red line is the growth in constant 40 °C, dashed lines and error bars show mean ± SE. Grey dashed line is predicted growth rate if there were no lag.

## Discussion

### Growth in fluctuating environments

We investigated stepwise fluctuations in temperature and could show that for *N. crassa* fast fluctuations slowed growth down more than expected based on growth rates under constant temperatures, and this was due to the time it took to recover from hot temperature. If fluctuations were faster than recovery times, the fungus never reached its higher growth rate in the cooler temperature. Once the duration of fluctuation steps increased beyond recovery time, growth in fluctuating environment sped up to or near the levels one would expect based on reaction norms in obtained in constant temperatures. The periods of the fluctuations used in this study were short and these kind of fluctutions may be rare in nature. However, the important question is how does the period of environmental fluctuations scale with acclimation time? In species where acclimation time is longer, environmental fluctuations with longer periods may have similar effects as reported here. This study used large shifts in temperature, that again may be rare in nature. It is therefore possible that allowing the cultures to first acclimate to intermediate temperatures first would change acclimation times. However, it is unlikely that results would change completely, for example growth rate at 35 ^°^C is higher than growth rate at 30 °C, and acclimation to stressfull temperature of 40 °C is nevertheless required.

### Role of epigenetic mechanisms in fluctuating environments

Our results suggest that epigenetic mechanisms have a small role in responding to fluctuating temperatures. This contrasts with our results of the role of epigenetic mechanisms in constant temperatures, where much stronger effects were observed on phenotypic plasticity (Kronholm *et al*., 2016). Possibly this is because of the time frame of environmental fluctuations investigated here may be faster than the normal timescale of epigenetic regulation, at least for the shortest step durations, which were well below estimated cell cycle times. Transcriptional response to environmental changes can be rapid (Causton *et al*., 2001; Chechik and Koller, 2009) and occur within minutes, and in yeasts the half-life of most proteins is around ten hours, but with considerable heterogeneity among different proteins (Christiano *et al*., 2014). While there is evidence that some chromatin state changes occur in a circadian manner (Sahar and Sassone-Corsi, 2013), and thus there was potential for epigenetic changes to occur during the fluctuations, not much is known about the time scale of epigenetic regulation in general.

We observed a rection norm change and slower acclimation for *set-1* mutant, which is deficient in histone 3 lysine 4 trimethylation, and has improper expression of certain genes (Raduwan *et al*., 2013). The H3K4me3 modification has been associated with 5’-regions of actively transcribed genes (Pokholok *et al*., 2005; Ardehali *et al*., 2011), and thus the absence of this histone mark may interfere with the proper activation of correct transcription response required for acclimation. The H3K4me3 mark is also required for proper expression of the circadian clock in *Neurospora* (Raduwan *et al*., 2013). The clock is affected by temperature (Crosthwaite and Heintzen, 2010), and temperature changes used in this study were large enough to reset the clock (Crosthwaite and Heintzen, 2010). However, in previous experiments mutants affecting the circadian period did not show any differences in growth rate (Gardner and Feldman, 1981). Thus, temperature mediated effects of *set-1* are unlikely to be mediated through the clock.

The histone mark histone 3 lysine 36 methylation, which is absent in the *set-2* mutant (Adhvaryu *et al*., 2005), may also be important in fluctuating environments. However, no acclimation effects were detected for *set-2*, so some other mechanism than acclimation time has to be in play. The role of H3K36me is likely to keep transcriptionally active regions of the chromosome in open conformation (Venkatesh *et al*., 2012) and the mark is present in actively transcribed regions (Hampsey and Reinberg, 2003; Morris *et al*., 2005). We have previously observed that *set-2* is important in temperatures above 25 °C (Kronholm *et al*., 2016). If the absence of H3K36 methylation slows or otherwise interferes with rates of transcription at high temperatures this can explain the obervation that the *set-2* mutant also affects growth in fluctuating environments.

It may be surprising that we did not observe significant effect for the *qde-2* mutant in fluctuating environments. In constant temperatures we have found evidence that *Neurospora* ARGONAUTE homolog encoded by *qde-2* is important in response to constant temperatures (Kronholm *et al*., 2016). We also observed this in the temperature shift experiments, where the *qde-2* mutant acclimated slower than the control strain. However, it may be that this difference is not large enough to be detected when we let the strains grow in fluctuating environments. QDE-2 protein is responsible for processing of diverse types of RNAs, such as siRNAs (Maiti *et al*., 2007), microRNA-like RNAs (Lee *et al*., 2010) in *Neurospora*. Small RNAs have been linked to temperature responses in different organisms (Chen *et al*., 2015; Kronholm *et al*., 2016; Fast *et al*., 2017), they have the potential to act as thermoregulatory molecules as they can adopt different conformations based on temperature, and especially in prokaryotes there are examples of RNAs acting as thermosensors (Sengupta and Garrity, 2013).

### Predicting growth in fluctuating environments

There has been lot of interest in predicting performance in fluctuating environments based on reaction norms measured in constant environments (Niehaus *et al*., 2012; Rezende *et al*., 2014; Ketola and Saarinen, 2015; Kingsolver *et al*., 2015; Kingsolver and Woods, 2016; Sinclair *et al*., 2016; Ketola and Kristensen, 2017). Our results show that prediction works in some circumstances: intuitively, if fluctuations are much slower than the time it takes to acclimate to a new temperature, then performance in constant environments predicts performance in a fluctuating environment. In addition, if fluctuations happen faster than acclimation time, then performance in fluctuating environments can be predicted to some degree if acclimation times, in our case lag functions, are known. However, there are some complications that may be general. Importantly, the lag functions we estimated for *N. crassa* were not symmetric and the shape of the functions are likely to change when the organism has only partially acclimated to a new temperature and the temperature changes again. This is reflected in our results, our refined lag model gave the worst predictions in step durations of 60 - 150 min where some acclimation had happened but which was not enough time to completely acclimate. With longer step durations predictions of the model were much better, and was nearly completely in line with observed data.

In a recent modeling study Kingsolver and Woods (2016) introduced a general model to predict performance in fluctuating environment specifically accounting for time-dependent effects. Time-dependent effects in this model were framed in terms of heat shock protein production and degradation. It was assumed that both production and degradation happen at similar rates, but this assumption is likely to be violated in several cases, as highlighted by asymmetric acclimation responses in our data. More empirical estimates of acclimation responses are needed so that these assumptions can be relaxed in future work on performance curves.

## Conclusions

Surprisingly, we did not find that epigenetic mechanisms are required for tolerating fluctuations that happen within one generation. There were some effects on acclimation time, but in these cases no memory effects seem to be involved, rather the physiological acclimation response seems to be impaired. Our results apply to within generation effects, and possible transgenerational effects need to be investigated separately. Predicting performance in fluctuating environments depends on the timescale of environmental fluctuations and acclimation functions of the organism in question. Increasing the complexity of the models allows taking lags into account but empirical data are needed to parametrize the models. An added complication is that acclimation seems to be asymmetric and the shape of acclimation functions is likely to change when the temperature range of acclimation changes. These results warrant caution when using reaction norms to predict performance in fluctuating environments. One cannot simply use data measured in constant environments and extrapolate to fluctuating environments, but by measuring acclimation functions with careful experiments, reasonably accurate models for growth in fluctuating environments can be constructed. Such experiments can also give insight into the mechanisms of acclimation.

## Acknowledgements

We’d like to thank the Fungal Genetics Stock Center (Manhattan, Kansas), Eric Selker, and Tereza Ormsby for kindly providing *Neurospora* strains, and Matthieu Bruneaux for critical reading of the manuscript. We’d also like to thank three anonymous reviewers for their constructive comments. This research was supported by the Academy of Finland (grants no. 274769 to I.K. and no. 278751 to T.K.), and the Centre of Excellence in Biological Interactions of University of Jyväskylä.

## Authors’ contributions

I.K. and T.K. conceived the study and designed the experiments, I.K. performed the experiments, I.K. analysed the data with input from T.K., I.K. wrote the manuscript with input from T.K. Both authors gave their final approval for publication.

## Data Archiving

Data and R scripts for analysis will be deposited in Dryad Digital Repository upon acceptance.

## Supplementary Information

**Figure S1:**
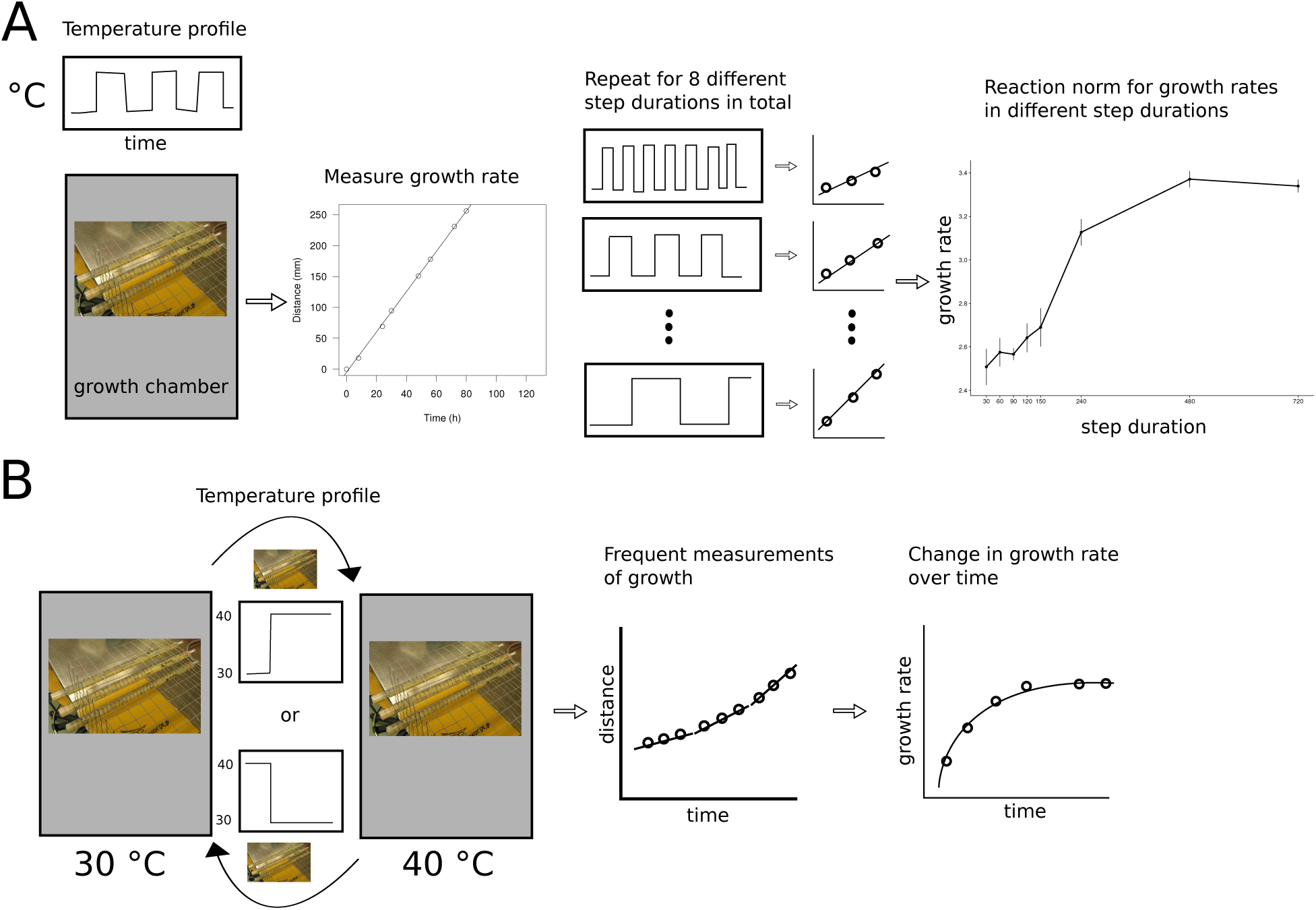
Overview of the experimental design. A) Obtaining growth rates for different fluctuating environments. Culture tubes were grown in a growth chamber where temperature fluctuated over time with a particular step duration. Measurements were taken to obtain growth rate for each tube. Growth rates were obtained for different step durations and from this data a reaction norm of growth rate for different step durations could be constructed. B) Temperature shift experiments. Two growth chamber compartments were set at 30 and 40 ^°^C. Temperature in the growth chambers was held constant, but the culture tubes were swapped between chambers. After the switch, measurements were taken at frequent intervals. From this data a profile of change in growth rate over time could be obtained.

**Figure S2:**
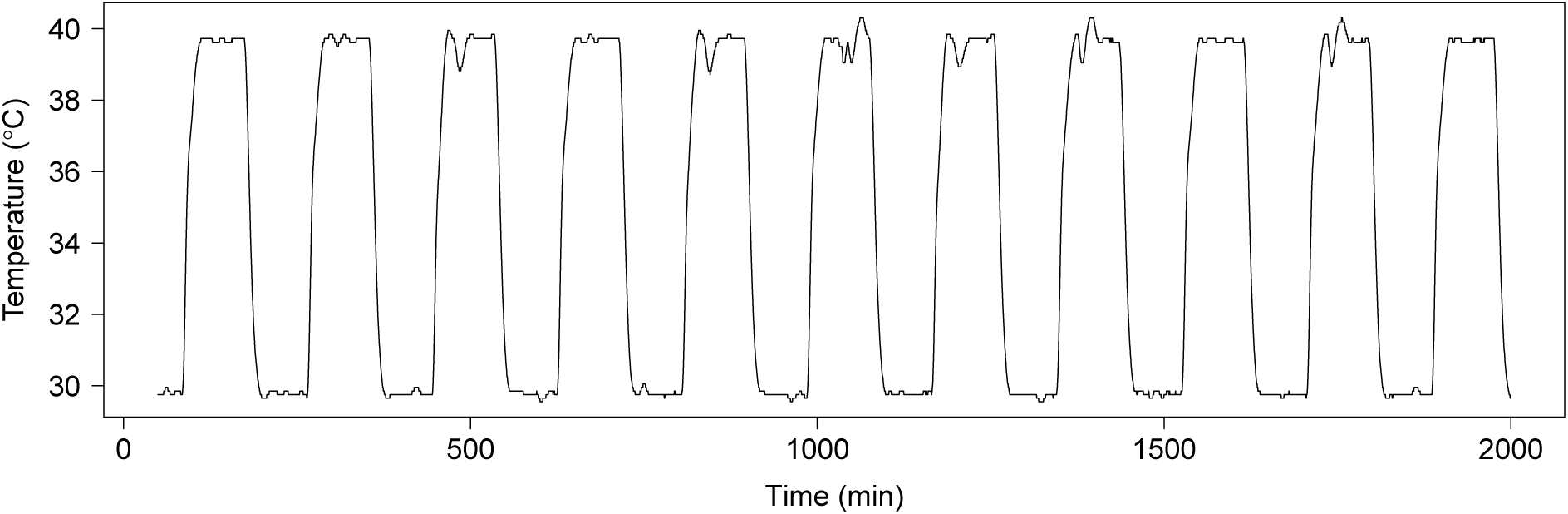
Realized temperatures from the experiment measuring growth rate in fluctuating environments. Temperature was recorded every 30 s with a datalogger. Step duration is 90 min in this example.

**Figure S3:**
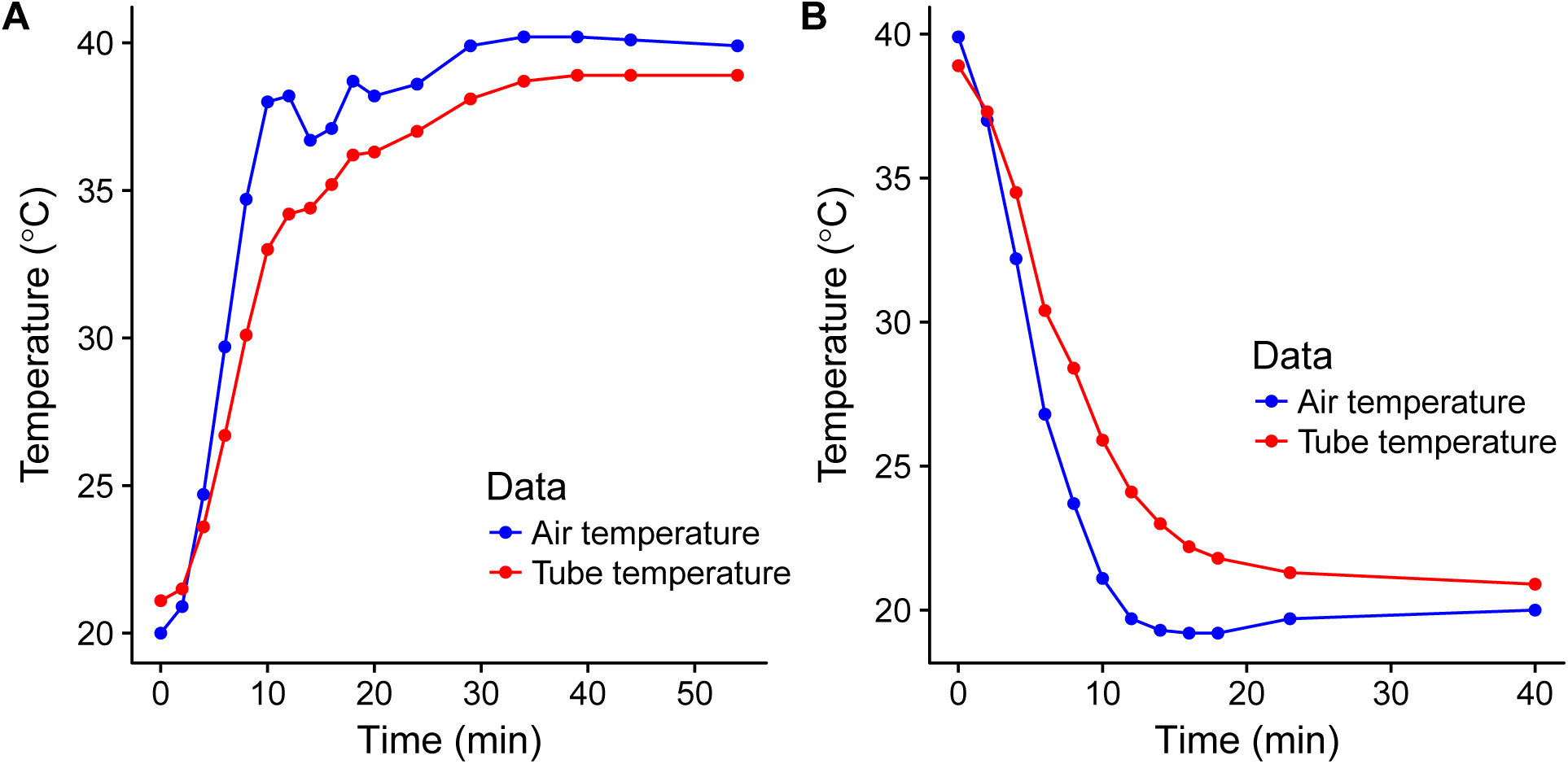
Changes in air and growth tube temperatures during growth chamber heating and cooling. In this preliminary experiment we measured changes from 20 to 40 ^°^C, to explore how fast temperature equilibrates between air and medium in maximal fluctuation. In the experiment temperatures altered between 30 to 40 °C. A) Growth chamber heating. B) Growth chamber cooling.

**Figure S4:**
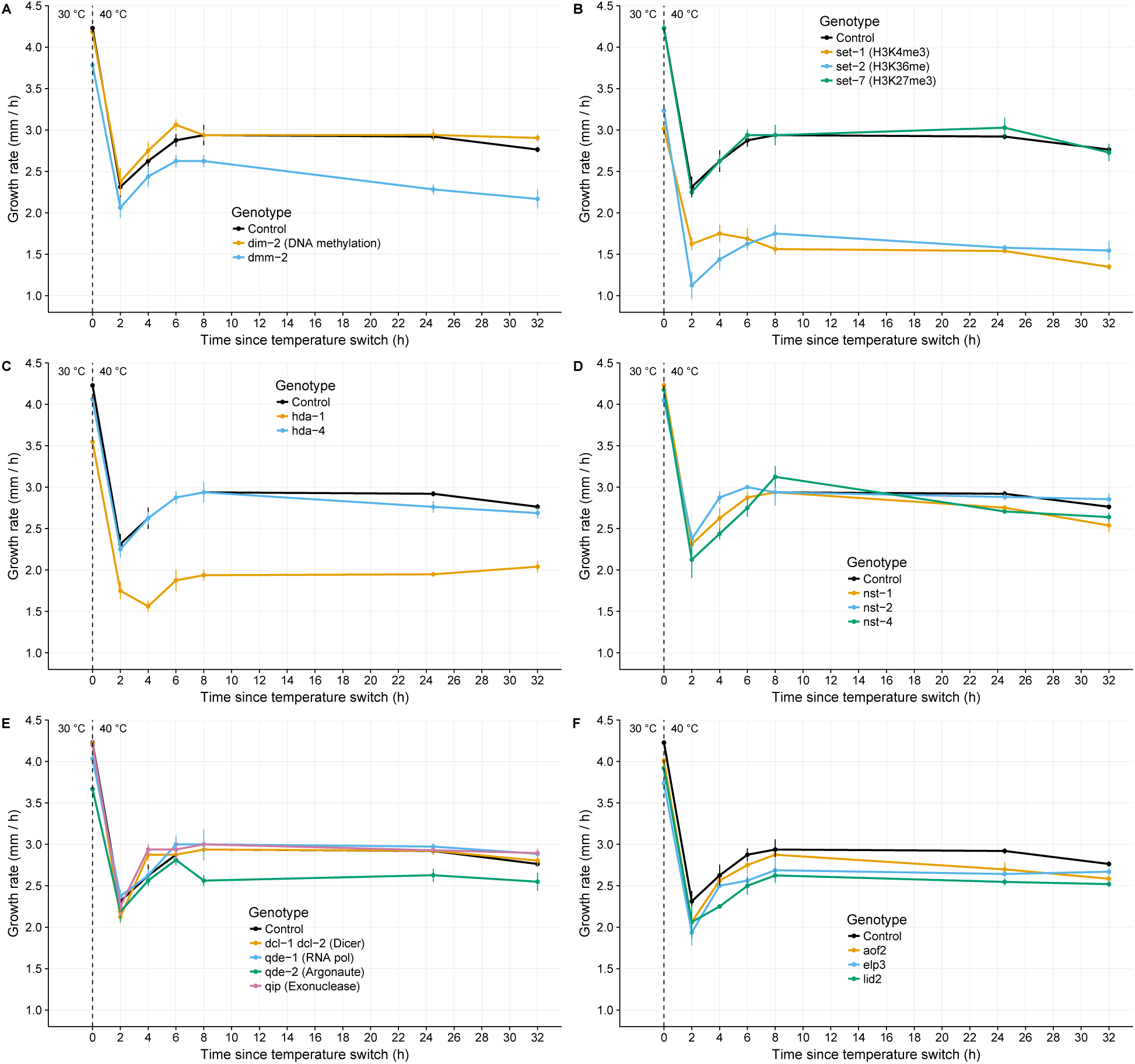
Changes in growth rate after temperature shift from 30 to 40 °C. Lines and error bars show means ± SE, n = 4. A) DNA methylation mutants. B) Histone methylation mutants. C) Histone deacetylation class I mutants. D) Histone deacetylation class III mutants. E) RNA interference mutants. F) Histone demethylation and acetylation mutants.

**Figure S5:**
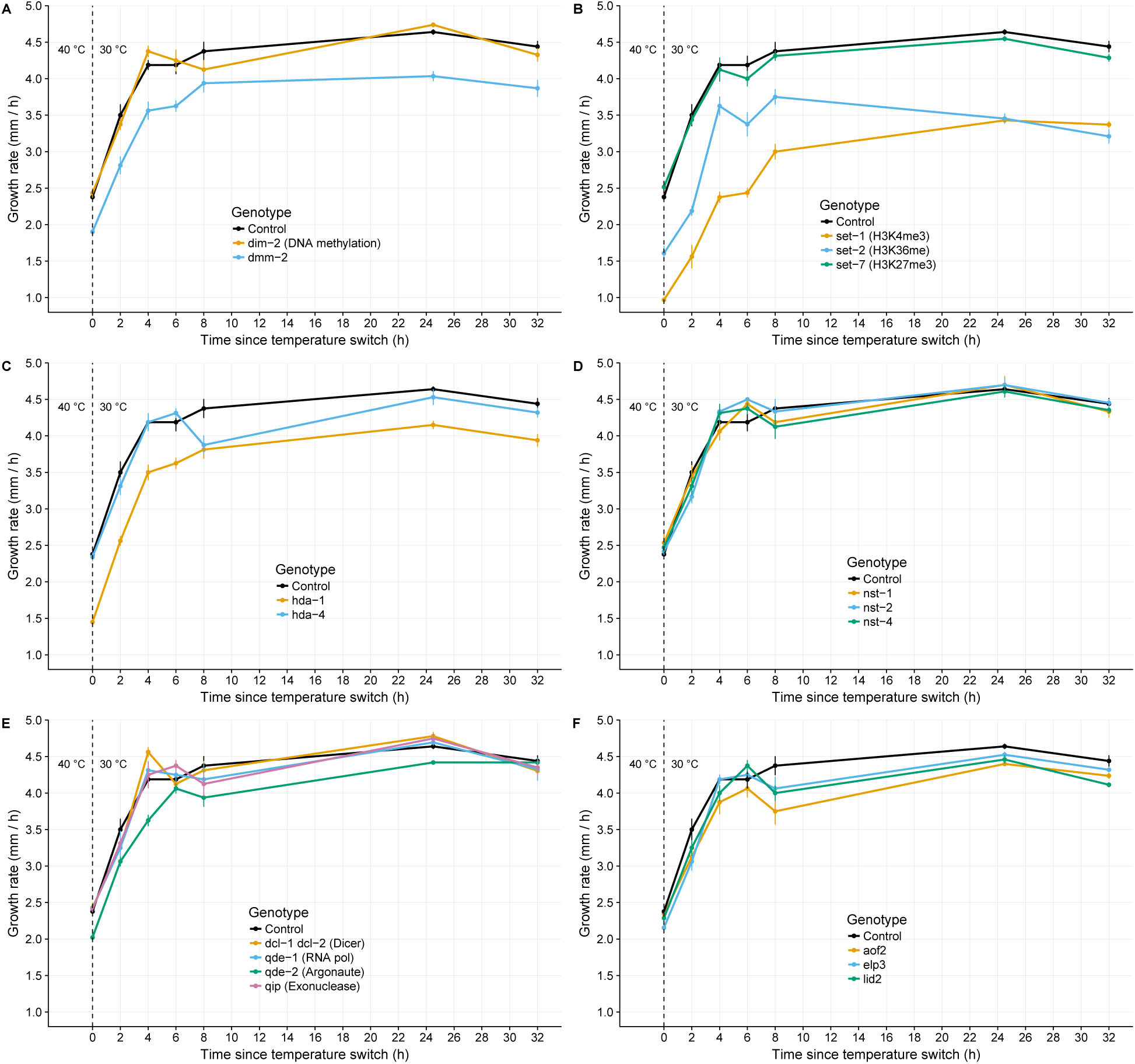
Changes in growth rate after temperature shift from 40 to 30 °C. Lines and error bars show means ± SE, n = 4. A) DNA methylation mutants. B) Histone methylation mutants. C) Histone deacetylation class I mutants. D) Histone deacetylation class III mutants. E) RNA interference mutants. F) Histone demethylation and acetylation mutants.

